# Large-scale RNA-seq mining reveals ciclopirox triggers TDP-43 cryptic exons

**DOI:** 10.1101/2024.03.27.587011

**Authors:** Irika R. Sinha, Parker S. Sandal, Grace D. Burns, Aswathy Peethambaran Mallika, Katherine E. Irwin, Anna Lourdes F. Cruz, Vania Wang, Josué Llamas Rodríguez, Philip C. Wong, Jonathan P. Ling

## Abstract

Nuclear clearance and cytoplasmic aggregation of TDP-43 in neurons, initially identified in ALS-FTD, are hallmark pathological features observed across a spectrum of neurodegenerative diseases. We previously found that TDP-43 loss-of-function leads to the transcriptome-wide inclusion of deleterious cryptic exons in brains and biofluids post-mortem as well as during the presymptomatic stage of ALS-FTD, but upstream mechanisms that lead to TDP-43 dysregulation remain unclear. Here, we developed a web-based resource (SnapMine) to determine the levels of TDP-43 cryptic exon inclusion across hundreds of thousands of publicly available RNA sequencing datasets. We established cryptic exon inclusion across a variety of human cells and tissues to provide ground truth references for future studies on TDP-43 dysregulation. We then explored studies that were entirely unrelated to TDP-43 or neurodegeneration and found that ciclopirox olamine (CPX), an FDA-approved antifungal, can trigger the inclusion of TDP-43-associated cryptic exons in a variety of mouse and human primary cells. CPX induction of cryptic exon occurs via heavy metal toxicity and oxidative stress, suggesting that similar vulnerabilities could play a role in neurodegeneration. Our work demonstrates how diverse datasets can be linked through common biological features and underscores that public archives of sequencing data represent a vastly underutilized resource with tremendous potential for uncovering novel insights into complex biological mechanisms and diseases.

## Introduction

RNA splicing is an essential process in eukaryotic cells that maintains the integrity of the transcriptome and proteome. The spliceosome plays a pivotal role in this process of allowing a single gene to give rise to multiple distinct mRNAs by precisely joining different combinations of exons during transcription. The resulting protein isoforms often exhibit tissue- or developmental stage-specific expression patterns, highlighting the role of alternative splicing in regulating gene expression and protein structure (*1–7*). Defects in RNA splicing can lead to significant alterations in the transcriptome and proteome which could contribute to cellular dysfunction and disease states (*8–10*). Certain RNA binding proteins (RBPs) help maintain transcriptome fidelity by acting as intronic splicing repressors. These splicing repressor RBPs recognize and bind to specific motifs located near aberrant splicing sites such as cryptic exons, thus repressing their incorporation into the mature mRNA during the splicing process (*11–20*).

TDP-43 performs its role in cryptic exon repression by preferentially binding to uracil-guanine repeats ([UG]_n_) in RNA (*11*, *21*, *22*). Inclusion of cryptic exons often leads to premature termination codons and nonsense-mediated decay (*23*), while cryptic exons in the 5’ or 3’ untranslated regions may impact processes such as mRNA stability and translation efficiency (*24*). Although TDP-43 may have multiple functions, recent studies have converged on the importance of cryptic exon repression in neurodegenerative disorders, particularly amyotrophic lateral sclerosis (ALS) and frontotemporal dementia (FTD). Over 97% of ALS and 45% of FTD cases exhibit TDP-43 pathology (*23*, *25–27*), which is characterized by nuclear clearance and cytoplasmic aggregation of TDP-43 protein. Recent studies have also associated TDP-43 pathology with other neurodegenerative disorders, such as Alzheimer’s disease and Huntington’s disease (*28–40*). While significant research has focused on the cytoplasmic aggregation of TDP-43, the nuclear clearance of the protein has emerged as a critical aspect of its pathophysiological role. Nuclear loss of TDP-43 leads to the inclusion of nonconserved cryptic exons (*11*, *17*, *22*, *23*, *41–43*), some of which are unique to specific cell types (*41*). These cryptic exons can be identified in both experimental models and patient samples using RNA-sequencing (RNA-seq) technologies.

While the majority of cryptic exons are out-of-frame, approximately 3% of TDP-43 cryptic exons introduce in-frame insertions into mRNA coding sequences (*41*). For example, the translated human cryptic exon in *HDGFL2* produces a neoepitope that can be detected in patients pre-symptomatically and potentially be used as a fluid biomarker for TDP-43 loss-of-function (*44–46*). The feasibility, however, of using cryptic exon-encoded peptides as biomarkers relies on their exclusive inclusion in affected tissues. Establishing baseline levels of cryptic exon inclusion across human tissues can be challenging to accomplish computationally without pre-processed resources tailored for this purpose. For instance, while the Genotype-Tissue Expression (GTEx) project provides a wealth of RNA-seq data across multiple human tissues, the official GTEx portal does not offer a straightforward way to perform custom splicing queries on a sample-by-sample basis. This limitation hinders the user’s ability to establish comprehensive baseline splicing profiles and assess the specificity of cryptic exon expression.

Previous efforts have been made to compile and process large-scale RNA-seq datasets to provide resources to investigate splicing patterns in samples, such as Snaptron (*47*) and Recount3 (*48*). These databases offer pre-processed RNA-seq data that can be mined to extract information on cryptic exon inclusion levels. However, the ability to effectively utilize these tools requires a certain level of bioinformatics expertise, making them less accessible to many biologists. To address this challenge, we have developed a user-friendly, web-based resource (SnapMine) that streamlines the process of querying Snaptron and extracting relevant splicing information. Our web-based tool enables biologists to rapidly profile splicing and gene expression data across the public archive without the need for bioinformatics experience. SnapMine aims to facilitate exploring alternative splicing’s role in health and disease by providing a more accessible interface to query under-utilized RNA-seq resources.

SnapMine is available at https://irika.shinyapps.io/snapmine/ and can be used to mine human and mouse RNA-seq datasets from The Cancer Genome Atlas (TCGA), GTEx, and the Sequence Read Archive (SRA) (*49–51*). To demonstrate SnapMine’s utility, we employed TDP-43 cryptic exons as a representative splicing phenotype and mined these RNA-seq archives for their inclusion patterns. By leveraging SnapMine to comprehensively profile cryptic exon inclusion across hundreds of thousands of datasets, we established essential baseline levels in diverse non-disease tissues and cell types. This reference will support the development of robust biomarkers of TDP-43 loss-of-function and empower future studies on the role of cryptic exons in disease.

SnapMine makes it simple and intuitive to uncover hidden connections across disparate datasets. As a proof-of-principle, we harnessed SnapMine to explore the upstream processes that can regulate TDP-43 by identifying cryptic exons in studies that had no prior connection to TDP-43 or neurodegeneration. By detecting splicing patterns through this integrated and unbiased approach, SnapMine identified datasets that could shed light on potential disease pathways that contribute to the initiation or progression of TDP-43 pathology. This represents a shift from traditional methods that focus on mining pre-selected datasets already known for their relevance and significance. Instead, searches can be performed across diverse datasets to find hidden links that no other approach could reveal. SnapMine provides a new, effective way to understand complex biological processes by harnessing the underutilized potential of large-scale public data archives.

## Results

### Mining the Snaptron database for alternative splicing events of interest

The Snaptron database (*47*) includes compiled exon-exon splice junction information across samples from three human RNA-seq sample databases: SRA (*49*), GTEx (*50*), and TCGA (*51*). Mouse splice junction information has also been extracted from the SRA. We queried the known TDP-43-associated cryptic exon targets identified from human and mouse TDP-43 knockdown datasets (*11*, *52–55*). Junction counts were extracted from the Snaptron database and the percent spliced-in (PSI) ratio was calculated using counts from each sample (Fig. 1A). For each sample, the PSI value ranged between 0-100% depending on the rate of cryptic junction inclusion as compared to the canonical junction. Metadata regarding the samples, such as SRA accession number or tissue type, was also extracted to give context to alternative splicing events.

**Figure 1:**
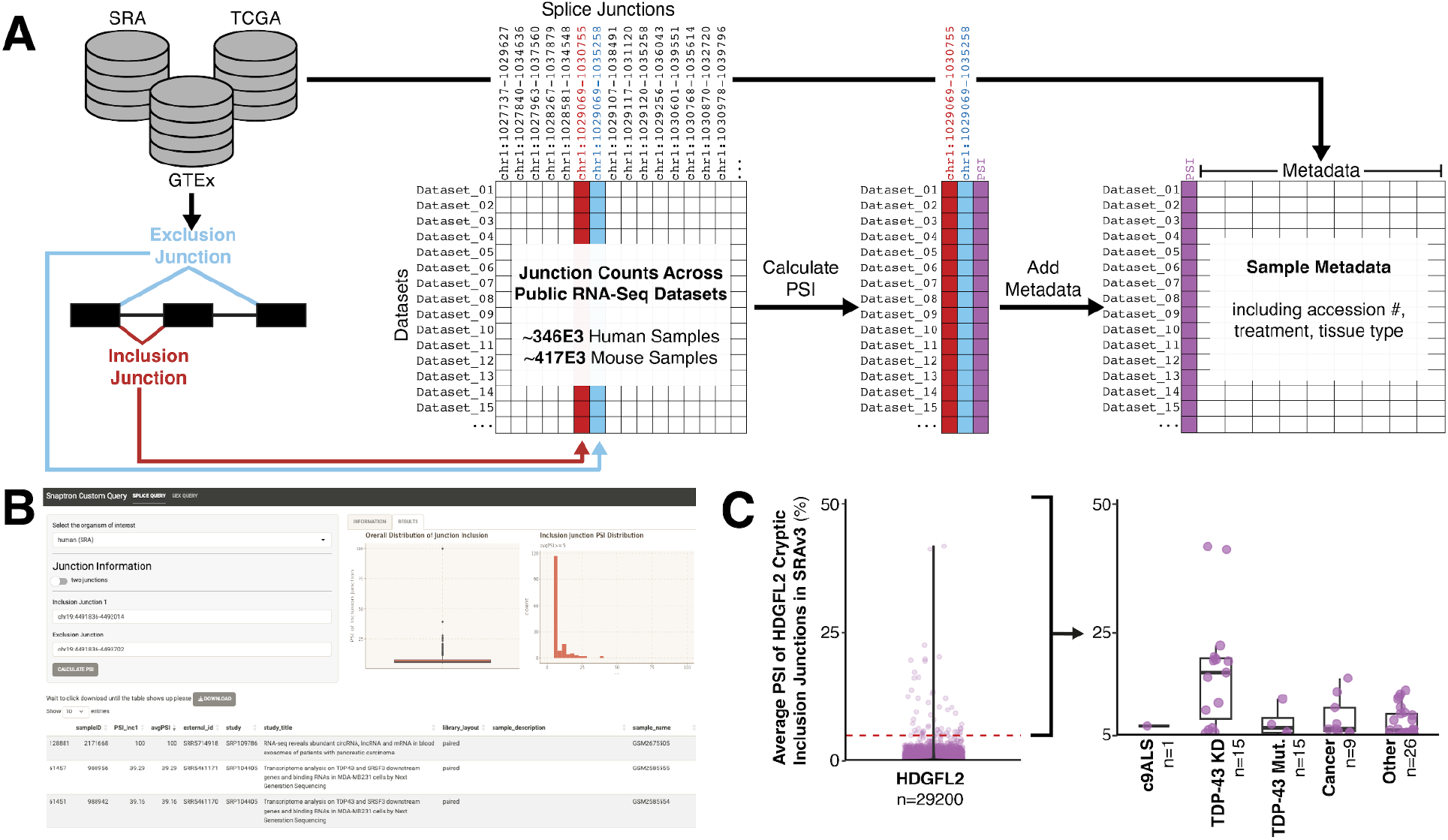
SnapMine application mines Snaptron database for exon-exon junction usage. (**A**) Schematic of SnapMine algorithm. (**B**) Example usage of SnapMine to identify one junction of the *HDGFL2* cryptic exon in the Sequencing Read Archive (SRA). (**C**) Only a few samples in the SRA include the *HDGFL2* cryptic exon at rates higher than 5%. Most of these are from ALS-associated or TDP-43-depletion associated studies.

R scripts were initially used to query Snaptron in singular and bulk for multiple cryptic exons. These scripts have been uploaded to GitHub and an R Shiny application to query databases for up to two junctions simultaneously has been made publicly available at https://irika.shinyapps.io/SnapMine (Fig. 1B). Users of the application can input inclusion and exclusion junction coordinates to calculate sample PSI values in one of the databases. The output of the query is a table detailing the PSI values of the inclusion junction for each sample with select metadata. If two junctions are queried, the average of the PSI values is also calculated. The table can be downloaded by the user and graphs of the data are generated and displayed for a quick visual regarding junction. Additionally, the application can tabulate gene expression counts within each sample using the pre-analyzed information in Snaptron. If desired, gene counts can be normalized to a control of choice. The example included in the application refers to *EDF1*, a gene with relatively constant levels of RNA expression levels across human tissues (*56*).

### HDGFL2 as a biomarker for TDP-43 loss of function

We first tested the capabilities of querying Snaptron by investigating the inclusion of the *HDGFL2* cryptic exon within the 316,000 human RNA-seq samples in the SRA compilation. We calculated the PSI for the junctions on the 5’ and 3’ sides of the cryptic exon compared to the canonical junction that skips the cryptic exon. In the 29,200 human samples in which the *HDGFL2* junction could be detected, only 55 samples (0.19%) had an average PSI for both cryptic junctions greater than 5% (Fig. 1C). Of the 55 samples, 18 were directly associated with TDP-43 depletion. Another was from a C9-ALS case, although the TDP-43 pathology level was not available. Finally, nine other samples were associated with cancer samples while the last 26 samples were not as easily categorized. Of uncategorized samples, conditions included viral/immune manipulations or other neurodegenerative disease. Overall, we found that the *HDGFL2* cryptic exon was included almost exclusively in cases of TDP-43 manipulation, dysregulation, or depletion and thus would be a relatively specific biomarker for cases of TDP-43 loss of function.

### Identifying baseline cryptic exon inclusion levels for potential biomarkers

Due to the recent interest in using TDP-43-associated cryptic exons as biomarkers, we also investigated the baseline PSI values for other identified cryptic exons. Of the many identified cryptic exons, we chose to test the application using those in *AGRN*, *ATG4B*, *G3BP1*, *HDGFL2*, *MYO18A*, *PFKP*, *RANBP1*, *STMN2*, *UNC13A*, and *UNC13B* due to their biomarker and therapeutic relevance (*44*, *46*, *52–55*, *57*, *58*). We queried these cryptic exons in the GTEx database, which includes RNA sequenced from many postmortem human organs generally expected to be free of disease processes (*59*) (Fig. 2A, Fig. S1A-B). For many of the cryptic exons, inclusion rates in these normal tissues are below 20% but the cryptic exon in *UNC13B* is a notable outlier. It is included at relatively high rates in most sequenced GTEx samples.

**Figure 2.**
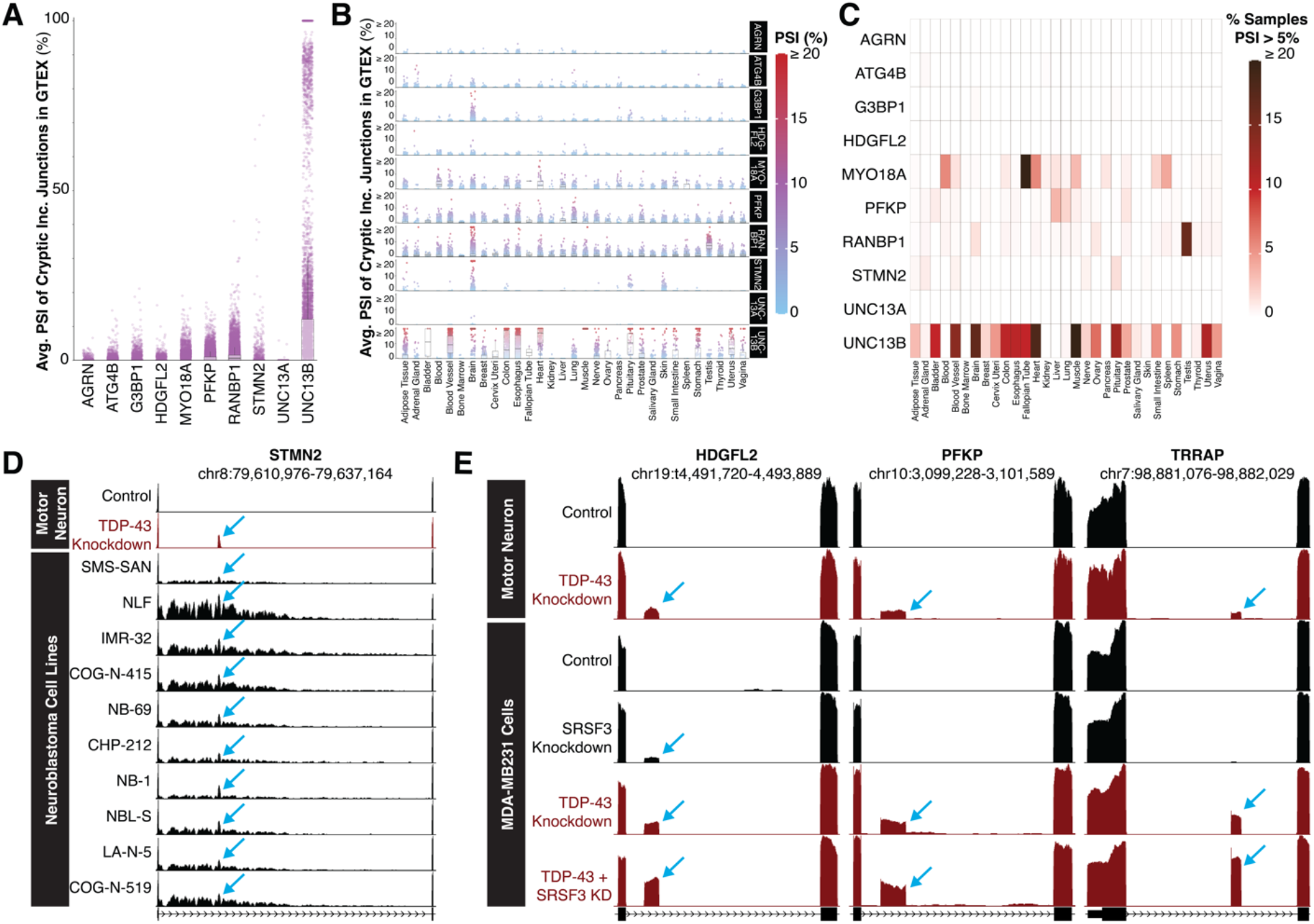
GTEx and SRA data compilations can be mined for cryptic exons of interest using SnapMine. (**A**) The average PSI of the cryptic exons in *AGRN, ATG4B, G3BP1, HDGFL2, MYO18A, PFKP, RANBP1, STMN2, UNC13A,* and *UNC13B* per sample is not always 0 in the GTEx dataset. The cryptic exon in *UNC13B* has particularly high inclusion rates. (**B**) Tissue-specific average PSI of each cryptic exon in GTEx samples. The average PSI of the *AGRN, ATG4B, G3BP1, HDGFL2, MYO18A, PFKP, RANBP1, STMN2, UNC13A,* and *UNC13B* cryptic exons differs by tissue sampled. Some cryptic exons have generally low inclusion in all samples while others have tissue-specific enrichment. (**C**) A heatmap representation of the percentage of samples from each tissue which have an average PSI of greater than 5% for the cryptic exon. *MYO18A*, *RANBP1*, and *UNC13B* have tissue-specific enrichment in GTEx samples compared to the other cryptic exons. (**D**) Visualization of the *STMN2* cryptic exon in some untreated neuroblastoma cell lines. (**E**) Visualization of MDA-MD-231 cells after TDP-43 and/or SRSF3 knockdown on UCSC Genome Browser (*62*). The PSI of *HDGFL2, PFKP,* and *TRRAP* cryptic exons appears to increase compared to a TDP-43 knockdown after co-knockdown of *SRSF3*. The *HDGFL2* cryptic exon is present after *SRSF3* knockdown only.

Due to the slight spread in PSI values for the cryptic exons, we investigated if there was any tissue-type specificity to the cryptic exons (Fig. 2B-C, Fig. S1A). The cryptic exons included at the lowest rates in the tested samples are those in *AGRN*, *ATG4B, HDGFL2,* and *UNC13A.* About 5% of the heart tissue samples indicate elevated levels of the *MYO18A* cryptic exon and certain brain samples exhibit higher splicing inclusion for cryptic exons in *RANBP1* and *STMN2*. This implies that TDP-43 loss-of-function may be more common in normal tissues than expected or that other mechanisms beyond TDP-43 dysfunction can contribute to cryptic exon incorporation (Fig 2C). Further work will be needed to identify conditions for cryptic exon inclusion prior to their usage as biomarkers in brain tissue or brain-related biofluids.

As a more comprehensive test of potential targets, we queried the inclusion of cryptic exons in GTEx for *CAMK2B*, *EPB41L4A*, *GPSM2*, *IGLON5*, *KALRN*, *PXDN*, *RSF1*, *SLC24A3*, *SYN1*, *SYT7*, and *TRRAP* (Fig. S1A, Table S1). PSI values of cryptic exons are generally low across most samples, but certain cryptic exons have elevated baseline splicing inclusion levels in certain tissues. Inclusion of the cryptic exon in *EPB41L4A* is slightly elevated in the esophagus, *PXDN* in the brain, *KALRN* in breast tissue, and *SLC24A3* in esophageal tissue. Overall, there are more samples with elevated PSI values for these TDP-43-associated cryptic exons in tissue derived from blood vessels, brain, colon, esophagus, heart, and muscle. Brain and muscle tissue show increased susceptibility to TDP-43-associated cryptic exon exclusion in GTEx tissue. It is important to note that the samples with high levels of TDP-43 associated cryptic exons may come from donors with undiagnosed TDP-43 pathologies such as ALS-FTD or inclusion body myositis (IBM). Additionally, there is no significant correlation between overall cryptic exon inclusion levels and age group in the GTEx samples (Fig. S1A-B).

### Querying SRA database for novel biological associations

After validating our ability to query databases through Snaptron for cryptic exon-associated junctions, we queried the SRA multiple cryptic exons associated with TDP-43 depletion to search for new associations or contexts in which the phenotype occurs. Unlike the GTEx database, the SRA contains numerous samples from tissues and cells that are not considered normal or have been subjected to environmental and/or genetic manipulations. Additionally, the databases indexed by Snaptron contain samples from both human and mouse sources. We mined the database for cryptic exons, removed the TDP-43-related samples, and searched for new associations.

First, we searched for the junction associated with the cryptic exon in *STMN2*, which leads to a polyadenylation event and subsequently reduces gene and protein expression (*40*, *52*, *58*, *60*). Current clinical trials are assessing a *STMN2*-targeting ASO for ALS treatment and the identification of non-ALS contexts of this cryptic exon inclusion could increase its potential therapeutic impact (*60*, *61*).

Our findings demonstrate expression of the *STMN2* cryptic exon in select neuroblastoma lines, including SMS-SAN, NLF, IMR-32, COG-N-415, NB-69, CHP-212, NB-1, NBL-S, LA-N-5, and COG-N-519 (Fig. 2D). While all these cell lines originate from pediatric neuroblastomas, there was little consistency regarding mutational profiles. For example, the NB-1 cell line has amplified *MYC* and *ALK* genes with wildtype levels of *p53* RNA while the COG-N-415 line also has amplified *MYC* and wildtype *p53* RNA levels but also has a F1174L mutation in *ALK*. Although inclusion levels vary, it is notable that *STMN2* cryptic exon is present in multiple lines. These results, in combination with the samples from the normal postmortem brain tissues which incorporated the *STMN2* cryptic exon (Fig. 2A-C), indicate that the expression of this cryptic exon is not limited to ALS-FTD and it may have leaky splice sites. Additionally, this excludes use of the *STMN2* cryptic exon as a measure of TDP-43 depletion in these neuroblastoma cell lines, and perhaps others, if the cell line is used as a model.

Next, we looked further into the inclusion of the cryptic exon in *HDGFL2* in SRA samples. As previously mentioned, 26 samples had a PSI of greater than 5% but were not easily categorized into samples associated with cancer, ALS, or TDP-43 (Fig. 1C). Surprisingly, multiple samples corresponded to a knockdown of Serine and Arginine Rich Splicing Factor 3 (SRSF3). These samples came from a dataset in which TDP-43 and SRSF3 were knocked down either singularly or in concert. In MDA-MB231 cells, the two proteins were found to form a complex and the loss of either caused splicing dysregulation (*62*). Analysis of the RNA-seq reads displayed the inclusion of multiple cryptic exons, including that in *HDGFL2*, after SRSF3 knockdown only (Fig. 2E). Additionally, a combinatorial knockout of both TDP-43 and SRSF3 led to increased incorporation of the cryptic exons in some genes, including *HDGFL2*, *PFKP*, and *TRRAP*, when compared to the single TDP-43 knockdown. Given SRSF3’s established role in alternative splicing (*63*), this supports the idea of the protein acting synergistically with TDP-43 to repress cryptic exons.

### Querying human SRA database for potential biomarker cryptic exon inclusion level

Following the success of querying the SRA database for *HDGFL2* and *STMN2* cryptic exons, we mined the database for the junctions associated with cryptic exons in *ACTL6B, AGRN, ATG4B, G3BP1, MYO18A, PFKP, RANBP1, UNC13A, and UNC13B* (Fig. 3A). For cassette cryptic exons, we averaged the PSI of both junctions in each sample. We found that the inclusion of cryptic exons with a PSI above 5% occurred in less than 1% of samples for *ACTL6B, AGRN, ATG4B, G3BP1, PFKP, RANBP1,* and *UNC13A.* This indicates that, generally, these cryptic exons have not been detected within the samples stored in SRA, whereas cryptic exons in *MYO18A, STMN2,* and *UNC13B* were included with a PSI greater than 5% in more than 1% of samples, potentially indicating greater leakiness. This confirms the results from the GTEx database that also show greater rates of inclusion in normal tissue (Fig. 2B, Fig S1B). Cryptic exons in *STMN2* and *UNC13B* are included at greater rates than the other cryptic exons as they had a PSI of greater than 15% in more than 1% of the queried samples. Unsurprisingly, the top 20 samples in the SRA that included high rates of cryptic exon inclusion across the different targets were associated with ALS-FTD tissue or TDP-43 knockdown (Fig. 3C, Table S2). Many of these samples were specifically identified to have nuclear clearance of TDP-43. This is consistent with the cryptic exons being repressed by TDP-43 normally and included when the protein is depleted or dysfunctional.

**Figure 3.**
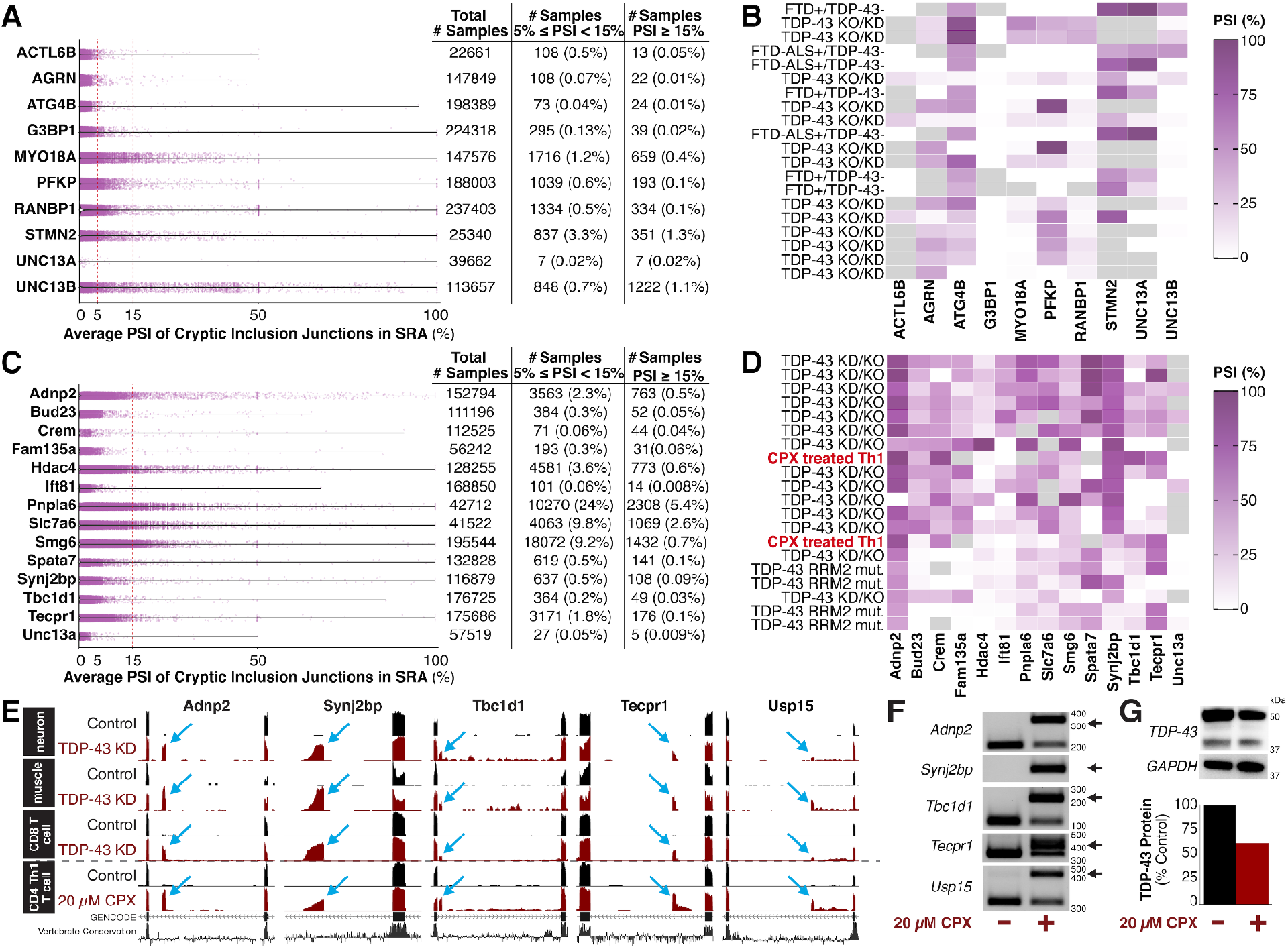
The SRA can be mined to identify novel biological contexts of alternative splicing events. (**A**) The average PSI of cryptic exons in *ACTL6B, AGRN, ATG4B, G3BP1, MYO18A, PFKP, RANBP1, STMN2, UNC13A,* and *UNC13B* can be extracted and quantified from human SRA datasets using SnapMine. Similar to the GTEx results, UNC13B is the most leaky event but some of the other cryptic exons also have greater than 5% inclusion in many samples. (**B**) The top 20 human samples from SRA with overall high inclusion of TDP-43-associated cryptic exons are associated with TDP-43 knockdown. TDP-43-samples are specifically depleted of nuclear TDP-43. (**C**) The average PSI of cryptic exons in *Adnp2, Bud23, Crem, Fam135a, Hdac4, Ift81, Pnpla6, Slc7a6, Smg6, Spata7, Synj2bp, Tbc1d1,* and *UNC13B* can be extracted and quantified from mouse SRA datasets using SnapMine. (**D**) Although the majority of the top 20 mouse samples from SRA with overall high inclusion of TDP-43-associated cryptic exons are associated with TDP-43 knockdown, two samples stand out. The two samples correspond to CD4+ Th1 cells treated with CPX *in vitro*. (**E**) CD4+ Th1 cells treated with CPX incorporate cryptic exons in *Adnp2, Synj2bp, Tbc1d1*, *Tecpr1*, and *Usp15* at levels comparable to mouse neuronal and muscle cells with TDP-43 knocked down. (**F**) Treatment of a mouse splenocyte culture with 20 µM CPX for four hours leads to cryptic exon inclusion as measured by RT-PCR. The black arrow points to the cryptic band. (**G**) Treatment of a mouse splenocyte culture with 20 µM CPX for four hours leads to TDP-43 depletion as measured by immunoblot. TDP-43 protein levels are normalized to GAPDH protein levels.

### Querying mouse SRA database for potential biomarker cryptic exon inclusion level

TDP-43-associated cryptic exons are non-conserved across species and, as such, those identified in mice cannot be targeted in humans. Interestingly, TDP-43 has the same function of splicing repression in mice as it does in humans and still binds UG-rich motifs. Due to the ubiquitous expression of TDP-43 and conserved function, it is possible that upstream regulators of the protein are shared between species. Mouse cells are often subjected to genetic or environmental manipulations that would not be possible in human tissue. As a result, there are a greater variety of conditions available in mouse sequencing datasets. For this reason, we mined the mouse SRA for junctions associated with previously identified cryptic exons in *Adnp2, Bud23, Crem, Fam135a, Hdac4, Ift81, Pnpla6, Slc7a6, Smg6, Spata7, Synj2bp, Tbc1d1, Tecpr1,* and *Unc13a* (Fig. 3C). We found that the inclusion of cryptic exons with a PSI above 5% occurred in less than 1% of samples for the cryptic exons in *Bud23, Crem, Fam135a, Ift81, Spata7, Synj2bp, Tbc1d1,* and *Unc13a.* The inclusion of cryptic exons with a PSI of 15% occurred in more than 1% of samples for *Pnpla6* and *Slc7a6*, indicating leakiness and the potential for other regulators of the splicing events.

As in the human samples, the majority of the top 20 samples that include cryptic exons are associated with TDP-43 manipulations, but two samples stood out from the rest. Like the TDP-43 depleted samples, they contained multiple cryptic exons with high PSI values. Uniquely, they had nothing to do with neurodegeneration, TDP-43, or splicing. Instead, these samples came from a study which treated mouse CD4+ T helper 1 (Th1) T cells with ciclopirox olamine (CPX), an FDA-approved antifungal medication that has also been studied for its properties as an iron chelator (*64*, *65*) (Fig. 3D). After the treatment, multiple cryptic exons were spliced into the gene transcripts at exceptionally high rates compared to the control (Fig. 3E). These include the cryptic exons in *Adnp2, Synj2bp, Tbc1d1, Tecpr1,* and *Usp15,* which can also be identified in TDP-43 depleted mouse cells.

### CPX treatment leads to robust cryptic exon inclusion across species and tissue types

In the original study, Th1 T cells were isolated and activated prior to a four hour treatment with 20 µM CPX (*64*). To validate the treatment effect in immune cells, we simplified the experiment and treated a single-cell suspension generated from mouse splenocytes for four hours with 20 µM CPX. We then harvested the cells and confirmed cryptic exon inclusion using RT-PCR.

Our initial test of the treatment led to measurable inclusions of cryptic exons in mouse splenocyte transcripts of *Adnp2, Synj2bp, Tbc1d1, Tecpr1,* and *Usp15* (Fig. 3F). Using protein immunoblot, we then confirmed that CPX treatment led to a protein level depletion of TDP-43 in the samples (Fig. 3G). This highlights the accuracy of querying the Snaptron databases and using SnapMine to find novel biological associations.

Due to the success in replicating cryptic exon inclusion after CPX treatment in splenocytes, we were also interested in whether TDP-43 protein dysregulation by CPX is a general phenomenon or was specific to mouse splenocytes. Since TDP-43 dysregulation in ALS-FTD occurs in the nervous system, we first expanded our experiments to the mouse brain. We treated a single cell suspension of the mouse brain with 20 µM CPX for four hours, similar to the splenocytes. After the treatment, we conducted RT-PCR using single-band primers directed towards the cryptic exons in *Ift81* and *Unc13a*, as well as double-band primers targeting *Camk1g*. We found that these cryptic exons are spliced into the mRNA transcripts after CPX treatment, indicating that TDP-43 is dysregulated (Fig. 4A). Additionally, treatment of a mouse brain in an *ex vivo* setting led to increased levels of cryptic exon inclusion in *Ift81* and *Unc13a* when measured using Basescope ISH assays (Fig. 4E), suggesting that the application of CPX *in vivo* could be used to trigger and model TDP-43 dysregulation.

**Figure 4.**
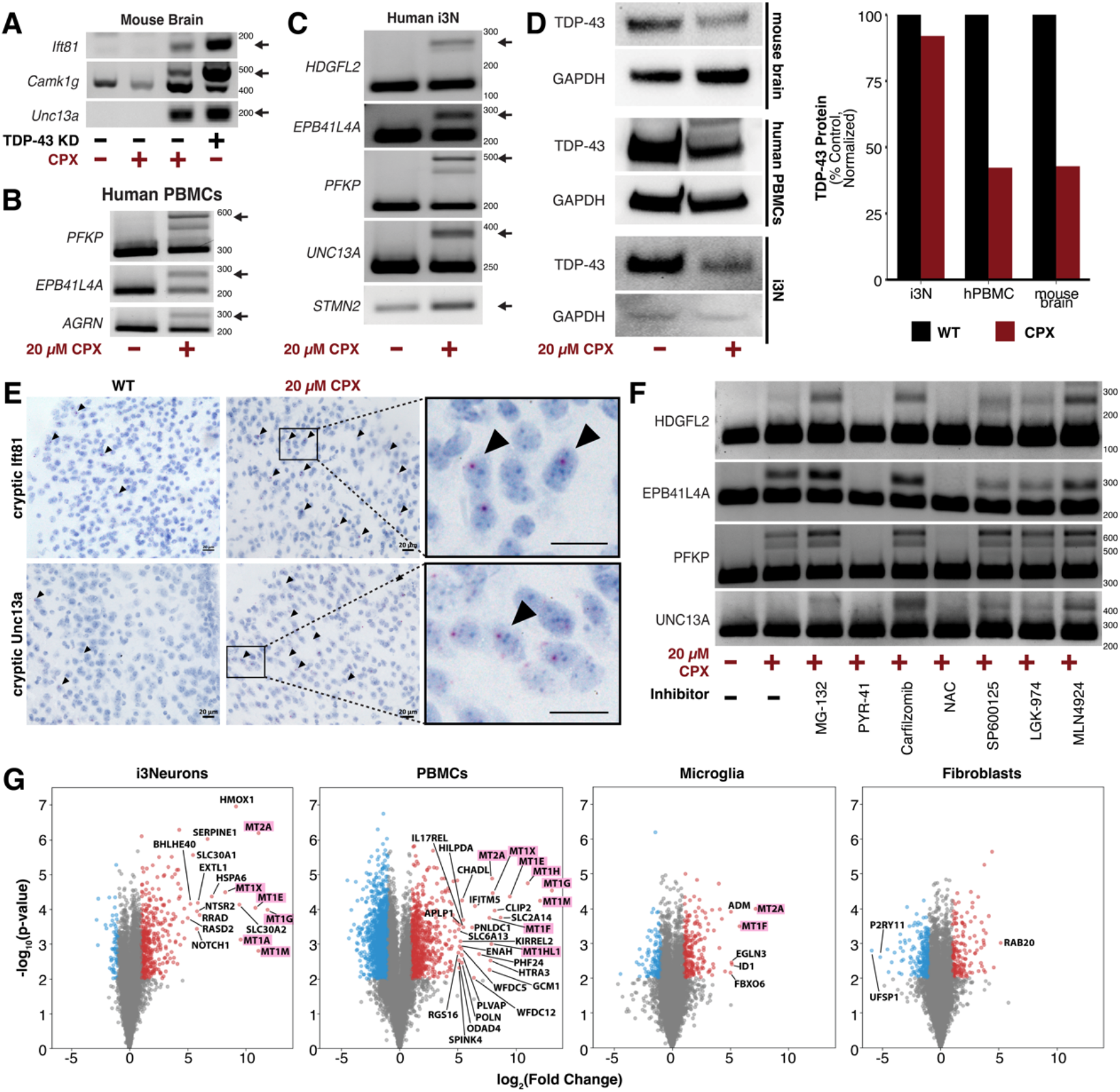
CPX treatment causes TDP-43 protein depletion in various primary cell types. (**A**) Treatment of a mouse brain culture with 20 µM CPX for four hours leads to cryptic exon inclusion as measured by RT-PCR. The black arrow points to the cryptic band. The primers used to measure *Ift81* and *Unc13a* cryptic exon inclusion target the cryptic exon directly and so no WT band is measured. (**B**) Treatment of a human PBMC culture with 20 µM CPX for four hours leads to cryptic exon inclusion in *PFKP, EBP41L4A,* and *AGRN* as measured by RT-PCR. The black arrow points to the cryptic band. (**C**) Treatment of a human i3N culture with 20 µM CPX for four hours leads to cryptic exon inclusion as measured by RT-PCR. The black arrow points to the cryptic band. (**D**) Treatment of a mouse brain, human PBMC, and human i3N cultures with 20 µM CPX for four hours leads to TDP-43 depletion as measured by immunoblot. TDP-43 protein levels are normalized to GAPDH protein levels. TDP-43 depletion varies by cell type. (**E**) *Ex vivo* treatment of mouse brain with 20 µM CPX for four hours leads to cryptic exon inclusion as measured by BaseScope probes targeting cryptic *Ift81* and *Unc13a*. (**F**) Treatment of a human i3N culture with 20 µM CPX for four hours in combination with different small compound inhibitors can change cryptic exon inclusion rates. Treatment with MG-132, Carfilzomib, and MLN4924 appear to increase cryptic exon inclusion rates consistently while treatment with NAC attenuates cryptic exon inclusion. (**G**) Volcano plots from RNA-Seq data comparing CPX-treated to untreated cells across i3Neurons, PBMCs, Microglia, and Fibroblasts (n=2 for each condition) show upregulation of distinct gene signatures that are specific to i3Neurons and PBMCs, cell types that exhibit cryptic exons after treatment with CPX. By contrast, microglia and fibroblasts do not exhibit cryptic exons after CPX treatment and likewise do not exhibit strong gene upregulation of metallothioneins (highlighted in magenta). Further analysis of genes upregulated in i3Neurons (Fig. S3) confirms that heat shock response and oxidative stress pathways are upregulated, suggesting that heavy metal toxicity may be mediating CPX’s effect on TDP-43.

We then expanded our search to investigate whether the mechanism of TDP-43 dysregulation by CPX is conserved in humans. Due to more robust incorporation of cryptic exons in the mouse spleen as compared to the brain, we decided to test samples containing human immune cells first and treated human peripheral blood monocytes (PBMCs) with 20 µM CPX for four hours. After the treatment, we found incorporation of cryptic exons in *PFKP*, *EPB41L4A,* and *AGRN* (Fig. 4B). This conservation of this effect across species led us to then to treat iPSC-derived neurons (i3N). We found that treated i3N incorporated cryptic exons, including those in *PFKP, HDGFL2, EPB41L4A, IGLON5, UNC13A,* and *STMN2* (Fig. 4C). Additionally, we found that the *STMN2* cryptic exon was sometimes incorporated at low levels in untreated i3N. These data indicated that the mechanism of TDP-43 dysregulation due to CPX treatment is not specific to mouse or immune cells and could be associated with pathways affected in ALS-FTD or other TDP-43 pathologies.

Although our results confirmed CPX’s effect leading to TDP-43 depletion in a cell-type specific manner (Fig. S2A), there were several potential mechanisms. During ALS-FTD, TDP-43 is normally depleted from the nucleus and sequestered in cytoplasmic aggregates. Immunoblot measurements of TDP-43 protein indicated CPX led to protein level depletion in these additional cell types although the extent of depletion varied by tissue type (Fig. 4D). These results indicated CPX treatment leads to increased degradation of TDP-43 protein.

Next, we aimed to determine the potential mechanism impacted by CPX, likely upstream of TDP-43 degradation. Initially, we pre-incubated mouse splenocyte samples with inhibitors of proteolysis or protein modification such as EST, a cysteine protease inhibitor, MG-132, a proteasome inhibitor, and PYR-41, a ubiquitin E1 inhibitor. We found that PYR-41 treatment reduced the level of cryptic exon inclusion, but further analysis found that it appears to have reduced the expression of many proteins in the cell, likely impacting many other pathways beyond TDP-43 (Fig. S2B). To test a more varied set of inhibitors in a human-relevant context, we pre-incubated the i3N with various inhibitors to determine whether TDP-43 function could be restored or diminished at a lesser level in the treated cells.

Using cryptic exon inclusion in *HDGFL2*, *EPB41L4A*, *PFKP*, and *UNC13A* as a readout of TDP-43 function, we found that certain inhibitors of the proteasomal degradation system, MG-132, Carfilzomib, and MLN4924, consistently led to an increase in the cryptic exon inclusion although they were not sufficient on their own to induce the cryptic exon (Fig. 4F, Fig. S2C). MG-132 and Carfilzomib directly inhibit the proteasome while MLN4924 impacts neddylation, another process involved in regulating proteasomal degradation of proteins (*66*). Additionally, pre-treatment with N-acetylcysteine (NAC), a reactive oxygen species (ROS) inhibitor, led to a noticeable decrease in cryptic exon inclusion. These results are consistent with an upstream mechanism of TDP-43 regulation that is partially dependent on proteasome and neddylation-associated degradation. Overall, our data suggests proteasome inhibitors, in conjunction with CPX, increase TDP-43 depletion, whereas, ROS inhibitors have the opposite effect, partially rescuing the CPX-induced depletion.

Finally, we performed RNA-Seq on i3Neurons, PBMCs, microglia, and fibroblasts to study gene expression changes between CPX-treated and untreated cells. Among these human primary cells, CPX induces cryptic exons only in i3Neurons and PBMCs, while microglia and fibroblasts appear to be resistant. Volcano plots generated from our RNA-Seq data (Fig. 4G) revealed significant upregulation of gene signatures specific to i3Neurons and PBMCs. Notably, these signatures did not appear in microglia and fibroblasts, which did not show cryptic exon inclusion after CPX treatment. Among the upregulated genes, metallothioneins were prominently upregulated both i3Neurons and PBMCs and a STRING network analysis (*67*) of genes upregulated in i3Neurons (Fig. S3) confirms the activation of heat shock response and oxidative stress pathways. These results suggest that cellular stress responses to heavy metals might be an underlying mechanism through which CPX exerts its effects on TDP-43, opening new avenues for understanding and potentially mitigating TDP-43 pathology in ALS-FTD and related disorders.

## Discussion

In this work, we have created an application that can be used to analyze thousands of samples in GTEx, TCGA, and SRA compilations for alternative splicing patterns of interest. The R script used to query for multiple cryptic exons simultaneously is available on GitHub and can be used by researchers interested in customizing the script. The application queries the Snaptron database of exon-exon junctions of interest and calculates the exon PSI values in each sample available. The output table includes metadata for each sample as annotated by Snaptron and can be downloaded as a csv file. Using the application, we were able to query the inclusion of TDP-43-associated cryptic exons in these databases and better understand baseline inclusions for consideration of each as a suitable biomarker.

To validate our application, we queried the inclusion of TDP-43-associated cryptic exons in these data compilations. GTEx includes biospecimens from donors collected within 24 hours of death and eligibility criteria is broad. As such, it is reasonable to expect that many of the donor organs will not be impacted by disease. Using this data, we determined that some cryptic exons can be found within organs without a TDP-43-associated deficit. Additionally, cryptic exon enrichment can differ on a tissue-to-tissue basis. Moving forward, the application can be used to determine baseline levels of cryptic exons and screen them in a tissue-specific manner for use as a biomarker or therapeutic target. By doing so, biomarkers specificity can be tested, and off-target effects of therapeutics can be avoided. Other researchers will also be able to use the application to mine public data for other novel biological contexts causing splicing abnormalities. Our preliminary analysis of the initial dataset of TDP-43-associated cryptic exon inclusion implicated CPX, an FDA-approved anti-fungal, as capable of causing TDP-43 depletion in a tissue-specific manner in as little as four hours. However, as the original dataset used CPX as an iron chelator, there were no explicit connections to TDP-43-associated pathology. Currently, CPX is available in prescription shampoo and cream formulations and off-brand uses include testing as an anti-cancer agent (*68–74*). Using the application, we were able to clearly identify TDP-43-associated cryptic exons in these CPX-treated samples and do further testing to explore what mechanism CPX may be targeting. Our in vitro experiments indicate the importance of caution when considering alternative applications of CPX, as it may impact proteins such as TDP-43.

Since we screened datasets using the average PSI value of multiple cryptic exons, there are likely other contexts in which single cryptic events occur. This provides the opportunity to find other upstream regulators of splice events. While we focused on TDP-43, it would be interesting to look at the targets of other RBPs, such as *PTBP1* or *MATRIN3* (*12*, *16*, *17*, *19*, *75–77*). Another potential future use would be to identify novel contexts of known alternative splicing and then re-analyze the RNA-seq data from implicated samples for new, context-specific RNA targets of the RBPs.

Our validation results with CPX also suggest that heavy metal toxicity and oxidative stress may be sufficient to induce TDP-43 dysregulation. Given that acute four hour treatments with CPX causes TDP-43 protein depletion, upstream mechanisms are likely related to protein modification or degradation process that occurs over short timespans. Additionally, we were able to show the process is conserved across multiple types of primary cell types beyond mouse Th1 T cells. Although not all cell types had TDP-43 dysregulation after 20 µM CPX treatment, its efficacy in human PBMCs and iPSC-derived neurons suggests that CPX treatment could be used in the future to create models of TDP-43 depletion. It is also possible that some cell types are more resistant to CPX-induced TDP-43 dysregulation and require higher levels or longer treatment.

Our work also supports prior research indicating increased TDP-43 dysfunction following proteasomal inhibition by MG-132 (*78*). While much of the previous research with TDP-43 and MG-132 focuses on ubiquitination and insolubility, 24 hour treatment with MG-132 also leads to the inclusion of cryptic *STMN2* in human motor neurons, which is indicative of nuclear depletion (*52*). While we did not find the four hour treatment of MG-132 sufficient for cryptic exon inclusion in i3Ns, combined treatment with both CPX and MG-132 led to increased cryptic exon inclusion when compared to only CPX. Supporting this hypothesis is the observation that treatment with Carfilzomib, a proteasome inhibitor used therapeutically, also increases cryptic exon inclusion when supplemented with CPX. This suggests proteasomal pathways play a role in TDP-43 degradation but are not the sole mechanism through which TDP-43 is degraded.

Since proteasomal inhibition led to worsened TDP-43 pathology, we hypothesize that CPX targets a different pathway which influences TDP-43 stability. If so, the combined treatment of CPX and proteasome inhibition would inhibit two different regulatory pathways of TDP-43 and increase its dysregulation. Previous treatment of cells with both niclosamide and a stressor found that treatment reduced levels of oxidative stress and TDP-43 pathology (*78*). Others have found that NAC treatment to reduce ROS has also reduced arsenite-induced stress granule formation and TDP-43 pathology (*79*). Our findings suggest CPX is another stressor that could lead to increased ROS, given that NAC treatment in combination with CPX partially alleviates cryptic exon burden. Some prior studies have also linked oxidative stress to TDP-43 aggregation and various neurodegenerative deficits (*80–82*). Thus, the observation that oxidative stress might contribute to nuclear clearance of the protein is compelling and points towards a common mechanism for TDP-43 pathology. Further investigation of the pathway through which CPX affects TDP-43 could elucidate information regarding endogenous regulators of TDP-43 and pave the way for the development of therapies that modulate these pathways in ALS-FTD and related dementias.

## Methods

### Analysis of RNA-seq Data

FASTQ files were downloaded from the NCBI’s Sequence Read Archive and aligned to the GRCh38 human genome assembly using STAR (v.2.7.10a) with default parameters (*83*). Megadepth was used to convert the output BAM files to BigWig files, and the data were visualized on the UCSC Genome Browser (http://genome.ucsc.edu/) (*84–86*). Alignment and conversion steps were completed on the Rockfish cluster at Johns Hopkins University.

### Analysis of Snaptron Data

All analysis was conducted using R version 4.3.2 on RStudio (R Core Team 2023). Inclusion and exclusion junction *hg38* or *mm10* coordinates were identified and then Snaptron junction counts for each sample were downloaded. PSI was calculated for each sample using the equation:

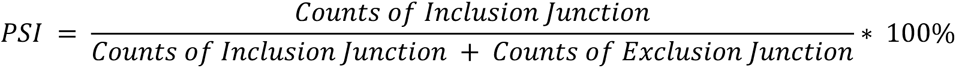

If the total number of junction counts, or the denominator of the above fraction, was less than 15, the sample was excluded from the analysis. Metadata downloaded from Snaptron was merged with the output PSI table. The *tidyverse* package was used to manipulate the data tables for analysis (*87*).

GEX counts were extracted for each sample from Snaptron using gene *hg38* or *mm10* gene coordinates and Ensembl gene ID of the queried gene. If normalization was desired, the counts were also extracted for the gene that is used for normalization. Relative quantitative gene count was calculated as follows. Normalization factor was calculated by determining the highest expression count of the normalization gene measured within a single study. For each sample, the normalization factor was the expression count of the normalization gene of the sample divided by the maximum value measured within the relevant study. The normalized count for the gene of interest was the gene expression count for the sample divided by the normalization factor for the sample. If there was no measurement of the normalization gene in a sample or study group, those sample(s) were marked as not normalized. This is depicted below.

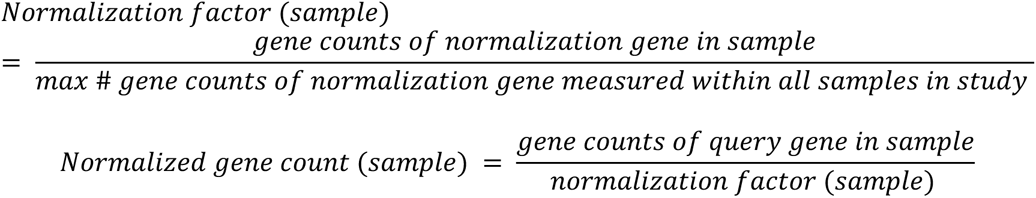

Visualization of Snaptron data was completed using the *ggplot2* and *viridis* packages in R (*88*). Final figures were made in Adobe Illustrator.

### R Shiny application and code availability

Usage of the above scripts for junction or gene expression count extraction and analysis from Snaptron are available at https://irika.shinyapps.io/SnapMine/ (*89*). Users can use the R Shiny application to query the TCGA, SRA, and GTEx for junctions of interest in available samples. The output table can be downloaded. The metadata files used to populate the table are static but can be supplemented using the original files on Snaptron. Scripts used to query junctions in bulk are available on GitHub (https://github.com/irikas/SnaptronMining). In addition to previously mentioned packages, the *ggthemr* package was used for visualization (github.com/Mikata-Project/ggthemr).

### Cell culture

To isolate mouse organs, the mouse was euthanized using carbon dioxide and perfused with PBS. Mouse spleen and brain were harvested as needed. To create a mouse splenocyte culture, the spleen was filtered through a 70 µm cell strainer, spun down, and red blood cells were removed using ACK Lysis buffer (Gibco #A1049201). Cells were resuspended in RPMI supplemented with 10% FBS and 1X Penicillin-Streptomycin (Gibco #15140122). To create a mouse brain cell culture, the brain was enzymatically dissociated (Miltenyi #130-107-677). Cells were resuspended in DMEM (Gibco #11960044) supplemented with 1X GlutaMAX (Gibco #35050061), 10% FBS, and 1X Penicillin-Streptomycin (Gibco #15140122). Cells were plated for half an hour and then treated.

The i3Neuron (i3N) iPSCs were cultured, grown, and differentiated as previously detailed (*90*, *91*). Neurons were treated on day 7 post-differentiation in cortical media. Human peripheral blood mononuclear cells (ATCC #PCS-800-011) were thawed as recommended and resuspended in RPMI 1640 medium (Gibco #11875093) with 1X GlutaMAX (Gibco #35050061), 10% FBS, and 1X Penicillin-Streptomycin (Gibco #15140122). They were plated for half an hour and then treated. Human dermal fibroblasts from healthy, male subjects from the Baltimore Longitudinal Study of Aging were cultured in DMEM, high glucose, HEPES medium (Gibco #12430054) with 10% FBS and 1X Penicillin-Streptomycin (Gibco #15140122). iPSC-derived microglia (Fujifilm iCell Microglia #01279) were thawed and cultured as per manufacturer recommendations. All cells were incubated at 37 °C and 5% CO2 during treatment and incubation/recovery steps.

### CPX treatment

Ciclopirox olamine, CPX, (Millipore Sigma #1134030-125MG) was solubilized in ethanol to 100 mM and 10 mM. Plated cells are treated with 20 µM CPX, or other concentration as indicated, for four hours in a 37°C tissue culture incubator. Cells were harvested after the treatment period and spun down. The supernatant was aspirated and the cells were flash frozen on dry ice for downstream analysis.

### Inhibitor treatment

Plated i3N were incubated with inhibitors at given concentrations for 30 minutes. After the pre-treatment period, cells were incubated with or without 20 µM CPX for four additional hours prior to harvest. The inhibitors, solvents, and concentrations tested are given below.

| Compound | Inhibits | Catalog Number | Solvent | Working Concentration |
| --- | --- | --- | --- | --- |
| 3-MA | Autophagy | Sigma-Aldrich 189490 | DMSO | 10 mM |
| Bortezomib | 20S proteasome | Sigma-Aldrich 5043140001 | DMSO | 100 nM |
| Carfilzomib | Proteasome | Selleck S2853 | DMSO | 35 nM |
| Compound C<br>(Dorsomorphin) | AMPK | Sigma-Aldrich 171261 | DMSO | 5 μM |
| LGK974 | Porcn Inhibitor<br>(Wnt pathway) | Sigma-Aldrich 5310910001 | DMSO | 1 μM |
| MG-132 | Proteasome | Sigma-Aldrich M7449 | DMSO | 10 μM |
| MLN4924 | Neddylation | Cell Signaling Tech 85923S | DMSO | 1 μM |
| N-acetylcysteine | ROS | Selleck S1623 | Water | 5 mM |
| PYR-41 | Ubiquitin E1 | Sigma-Aldrich 662105 | DMSO | 50 μM |
| SP600125 | JNK, Ser/Thr Kinase | Selleck S1460 | DMSO | 10 μM |
| Z-VAD-fmk | Pan caspase | Selleck S7023 | DMSO | 100 μM |

### RT-PCR analysis

RNA was extracted from samples using the Monarch® Total RNA Miniprep Kit (New England Biolabs, #T2010S) and cDNA was synthesized using the ProtoScript II First Strand cDNA Synthesis Kit (New England Biolabs, #E6560L). cDNA was amplified using primer pairs designed to either target two wildtype sequences on either side of the cryptic exon or one wildtype sequence and another sequence directly in the cryptic exon. Target sequence amplification was performed using Phusion Plus Green PCR Master Mix (Thermo Scientific, #F632L) and a modified touchdown PCR protocol (*92*), described below. Amplified targets were then separated by 1.5% agarose gel electrophoresis and visualized by ethidium bromide staining.

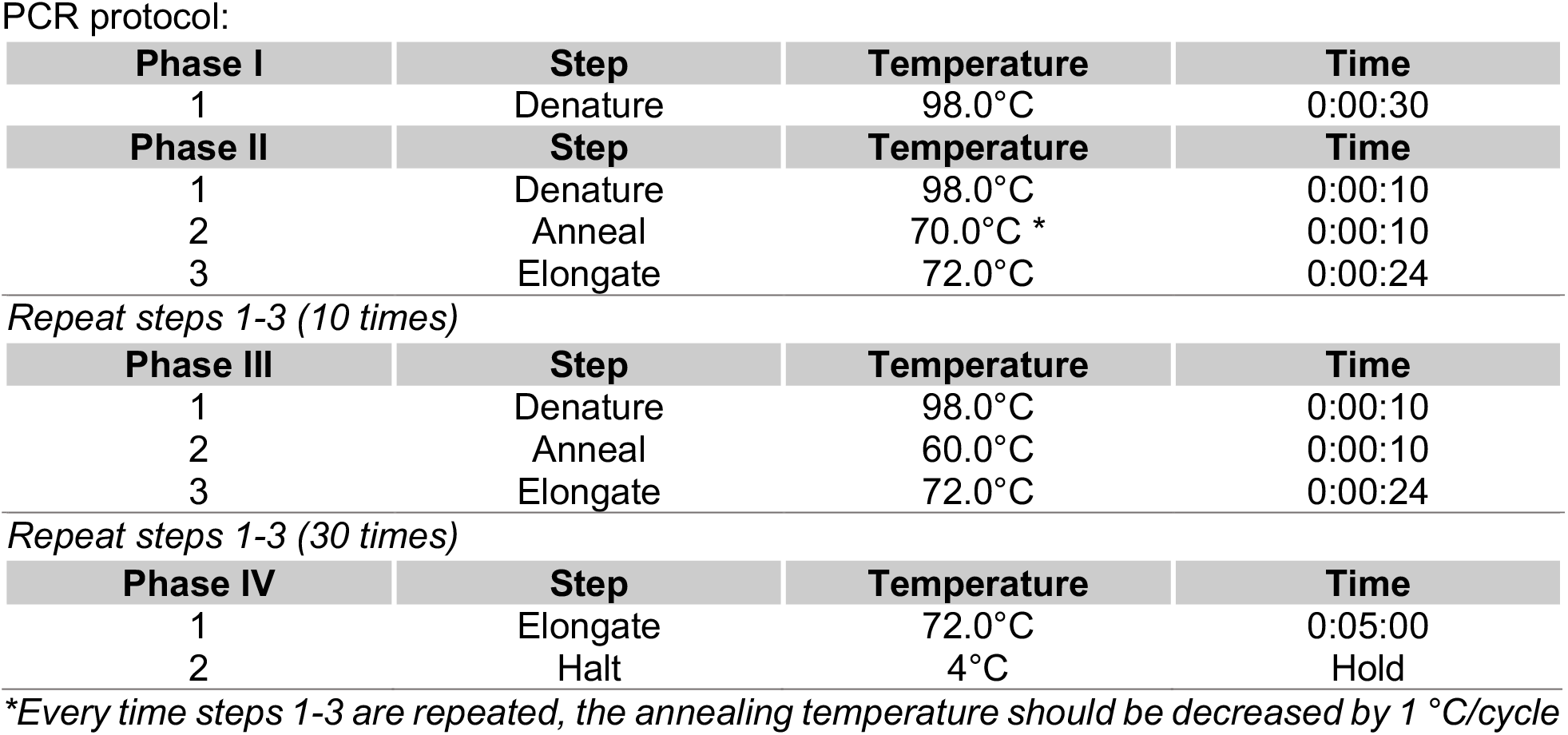

| Phase I | Step | Temperature | Time |
| --- | --- | --- | --- |
| 1 | Denature | 98.0°C | 0:00:30 |
| Phase II | Step | Temperature | Time |
| 1 | Denature | 98.0°C | 0:00:10 |
| 2 | Anneal | 70.0°C * | 0:00:10 |
| 3 | Elongate | 72.0°C | 0:00:24 |
| *Repeat steps 1-3 (10 times)* |  |  |  |
| Phase III | Step | Temperature | Time |
| 1 | Denature | 98.0°C | 0:00:10 |
| 2 | Anneal | 60.0°C | 0:00:10 |
| 3 | Elongate | 72.0°C | 0:00:24 |
| *Repeat steps 1-3 (30 times)* |  |  |  |
| Phase IV | Step | Temperature | Time |
| 1 | Elongate | 72.0°C | 0:05:00 |
| 2 | Halt | 4°C | Hold |

\*Every time steps 1-3 are repeated, the annealing temperature should be decreased by 1 °C/cycle

We used two types of primer pairs to amplify targets. The first type comprises a primer targeting the cryptic exon and another targeting an adjacent canonical exon, which we term single-band primers. Consequently, RT-PCR amplification would occur only when transcripts containing cryptic exons are in the sample. This leads to the amplification of cryptic exon-containing RNA at low levels. In the second type, both primers bind canonical mRNA flanking the cryptic exon, which we term double-band primers. Accordingly, two potential transcripts are amplified through RT-PCR: the wildtype and the heavier cryptic exon-including one.

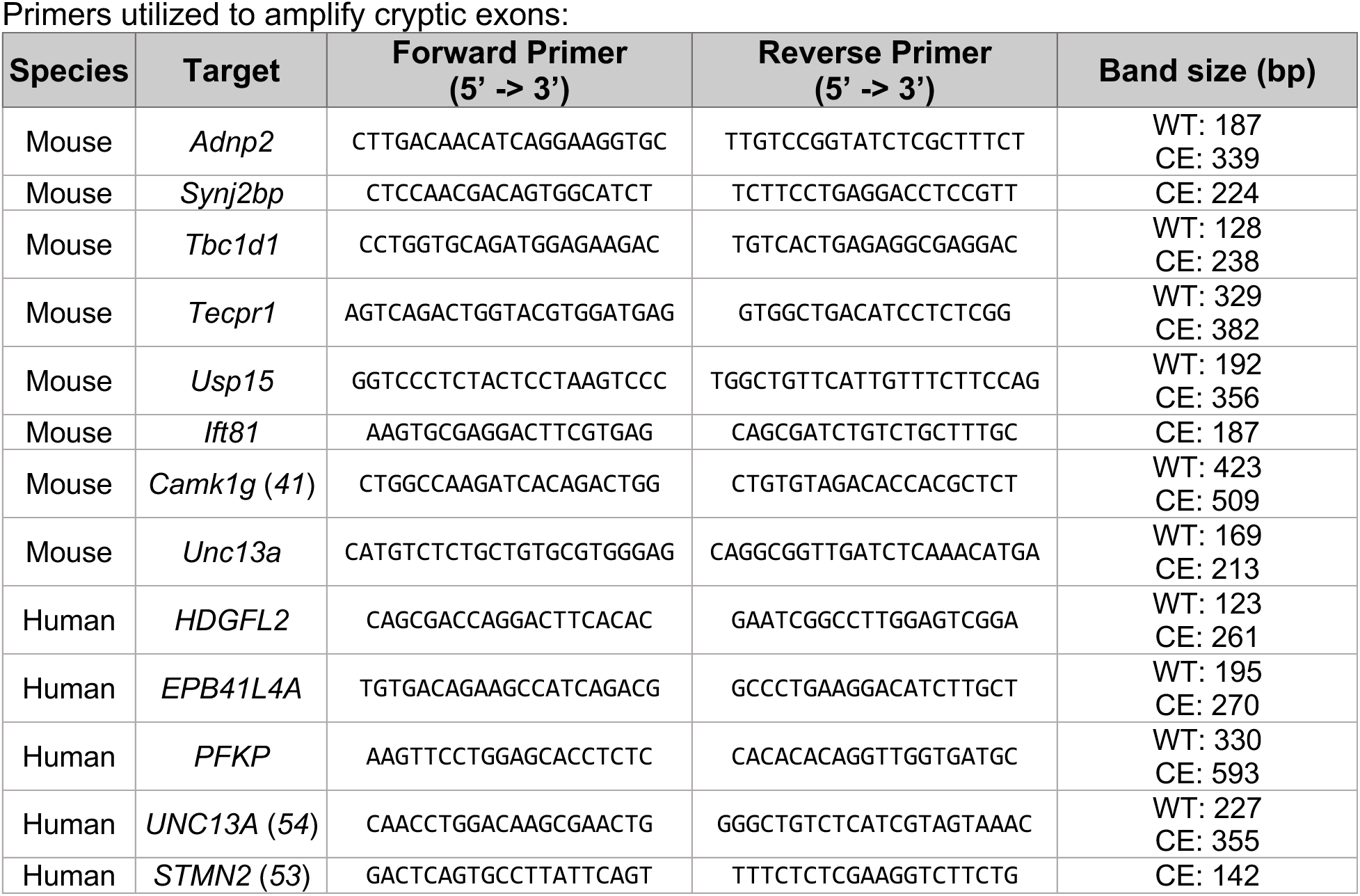
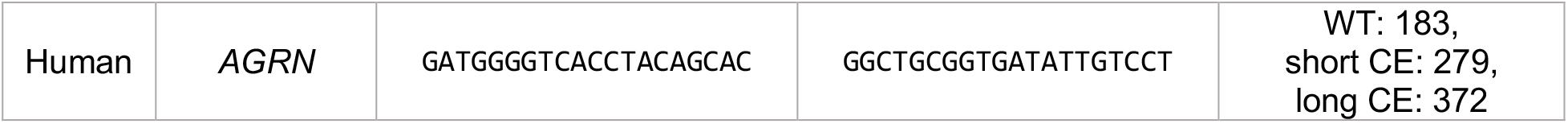

### Protein immunoblot and analysis

Samples were collected from wells by scaping and suspension in PBS. The cell suspension was centrifuged at 1600 x g for 5 min to pellet the cells. PBS was aspirated and the cell pellet was then resuspended in RIPA Lysis and Extraction Buffer (Thermo Scientific, #89900) with Halt Protease Inhibitor Cocktail (Thermo Scientific, #78430). After 5 minutes at 4C, the extract was centrifuged for 10 minutes at 4C and 16000 x g. The cell pellet was discarded and the Pierce BCA Protein Assay kit (Thermo Scientific, #23225) was used to quantify protein lysate concentration.

Protein blot analysis was performed following electrophoresis of reduced samples using NuPAGE 4 to 12% Bis-Tris Mini Polyacrylamide Gels (Thermo Scientific, #NP0322BOX). Protein was then transferred to PVDF membranes using the iBlot 2 Dry Blotting System from Invitrogen. The membrane was then used for western blot detection in one of two ways.

1. The membrane was then incubated in blocking buffer, 5% dry nonfat milk in Tris-buffered saline with 0.1% Tween (TBS-T). Other than anti-GAPDH primaries, membranes were probed with primary antibody in blocking buffer overnight at 4C with rocking. Secondary conjugated to HRP in blocking buffer was applied the next day for one hour at RT.
2. Alternatively, the membrane was pre-activated in methanol, soaked in iBind™ Flex Solution (ThermoScientific, #SLF2020), and probed using the iBind™ Flex Western Device (ThermoScientific, #SLF2000) according to manufacturer’s protocol

Immobilon® ECL UltraPlus Western HRP Substrate (Millipore Sigma, #WBULP-100ML) was used for detection in both cases. The BioRad ChemiDoc Imaging System was used to image the blots.To detect GAPDH, membranes were stripped using Restore™ PLUS Western Blot Stripping Buffer (Thermo Scientific, #46430) and reprobed using an anti-GAPDH antibody conjugated to HRP. Detection occurred as previously described.

| Target | Host | Company | Catalog # | Concentration |
| --- | --- | --- | --- | --- |
| TDP-43 | Rabbit | ProteinTech | 10782-2-AP | 1:2000 |
| TDP-43 | Rabbit | ProteinTech | 81350-1-RR | 1:5000 |
| GAPDH | Rabbit | ProteinTech | HRP-60004 | 1:5000 |

### BaseScope-ISH and Co-detection assay

RNA in-situ hybridization was performed using BaseScope Detection Reagent v2-RED Assay Kit (Advanced Cell Diagnostics, Inc. #323900), following manufacturer’s instructions. Expression of transcripts containing cryptic exon splice sites in *Unc13a* and *Ift81* transcripts was detected using 3zz custom BaseScope probes (BA-Mm-Unc13a-O1-2EJ-C and BA-Mm-Ift81-E17-intron17-NJ, respectively). Both positive and negative control probes were employed to assess the RNA quality (BA-Mm-Ppib and BA-DapB). Briefly, consecutive tissue sections of 10 μm thickness underwent de-paraffinization followed by pre-treatment with hydrogen peroxide, target retrieval buffer, and protease IV. Subsequently, these sections were subject to hybridization with target probes in a HybEZII oven (Advanced Cell Diagnostics, Inc.) for 2 hours at 40°C. The signals were amplified, and the slides were counterstained with hematoxylin. Images were acquired using a Zeiss Apotome Inverted Brightfield Microscope (Zeiss, Germany), and RNA puncta for each cryptic exon were manually quantified using ImageJ software.

## Acknowledgements

This work was supported in part by the Alzheimer’s Association (to J.P.L.), the Institute for Data-Intensive Engineering and Science (to J.P.L.), the NIH (nos. R01NS095969, UH3NS115608 and R33NS115161 to P.C.W.), the Robert Packard Center for ALS Research at Johns Hopkins (to P.C.W.), the Target ALS Foundation (to P.C.W.), ALS Finding a Cure (to P.C.W.), the ALS Association (to P.C.W.), the US Food and Drug Administration (no. 1U01FD008129 to P.C.W. and J.D.B.), and the NSF Graduate Research Fellowship Program under Grant No. DGE2139757 (to I.R.S.). Computational work was carried out at the Advanced Research Computing at Hopkins (ARCH) core facility (rockfish.jhu.edu), which is supported by the NSF (no. OAC 1920103).

## Data availability

Publicly deposited RNA-seq data in this study is available on the National Center for Biotechnology Information’s Sequence Read Archive under SRA study numbers SRP166282 (*52*), SRP057819 (*11*), SRP092413 (*93*), SRP104405 (*62*), SRP194266 (*94*), SRP367696 (*95*), and SRP079236 (*64*). NAUC data tables are available on http://ascot.cs.jhu.edu/ (*56*). The application to query junction inclusion data is hosted at https://irika.shinyapps.io/SnapMine.

## Contributions

I.R.S, P.C.W, and J.P.L. conceptualized the study. I.R.S and J.P.L wrote the manuscript, and all other authors edited and approved it. I.R.S., P.S.S., G.D.B, A.P.M, K.E.I., A.C, V.W., and J.L.R. performed experiments and/or analyzed data.

