## Supplementary Data for "Large-scale RNA-seq mining reveals ciclopirox triggers TDP-43 cryptic exons"

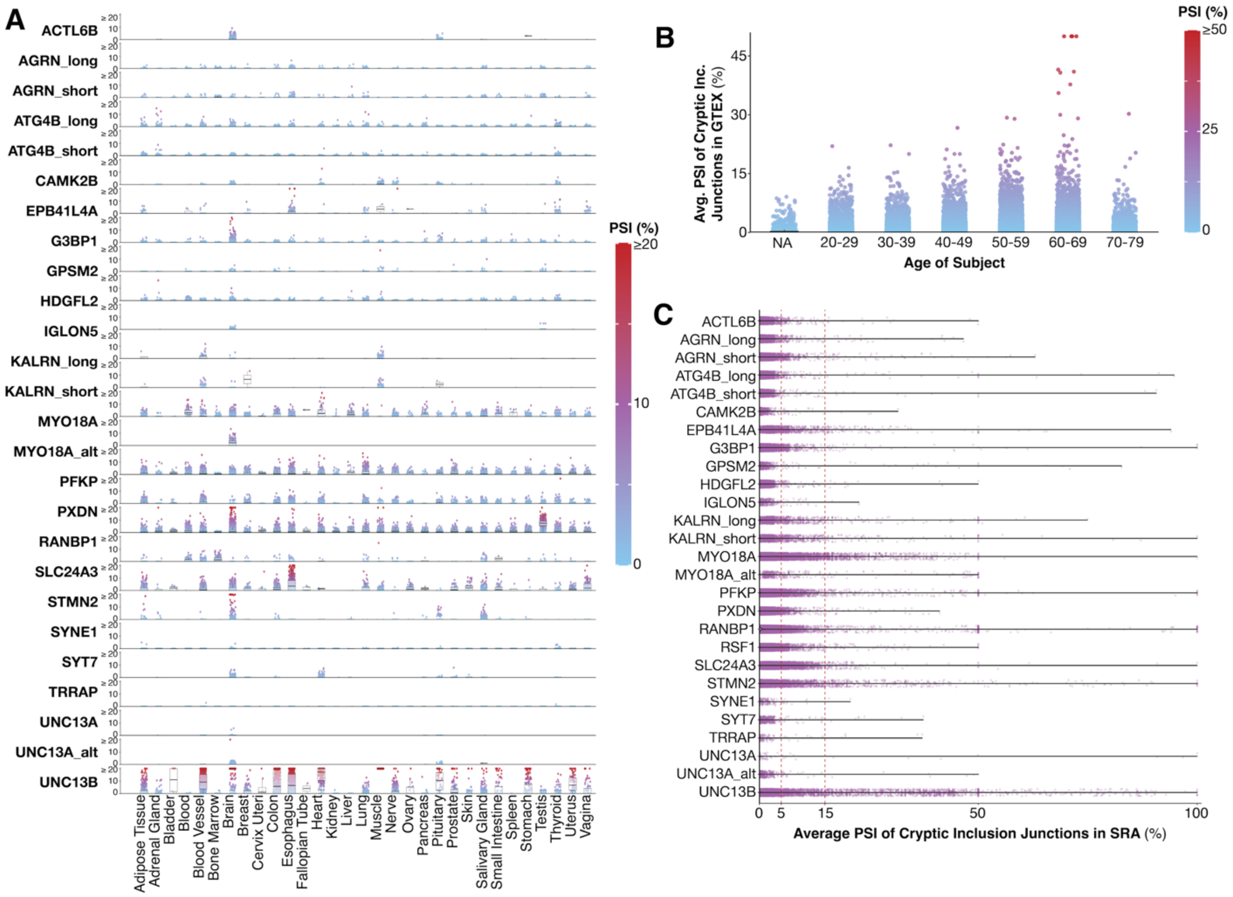


**Supplementary Figure 1: Using SnapMine to investigate GTEx and SRA data compilations can be mined for numerous additional cryptic exons.** (**A**) The average PSI of cryptic exons in *ACTL6B*, *AGRN*, *ATG4B*, *CAMK2B*, *EPB41L4A*, *G3BP1*, *GPSM2*, *HDGFL2*, *IGLON5*, *KALRN*, *MYO18A*, *PFKP*, *PXDN*, *RANBP1*, *SLC24A3*, *STMN2*, *SYNE1*, *SYT7*, *TRRAP*, *UNC13A*, and *UNC13B* can be extracted from and quantified in human GTEx datasets by tissue type using SnapMine. (**B**) Cryptic exon PSI values in human GTEx datasets by age. (**C**) The average PSI of cryptic exons in *ACTL6B*, *AGRN*, *ATG4B*, *CAMK2B*, *EPB41L4A*, *G3BP1*, *GPSM2*, *HDGFL2*, *IGLON5*, *KALRN*, *MYO18A*, *PFKP*, *PXDN*, *RANBP1*, *SLC24A3*, *STMN2*, *SYNE1*, *SYT7*, *TRRAP*, *UNC13A*, and *UNC13B* can be extracted from and quantified in human SRA datasets using SnapMine.

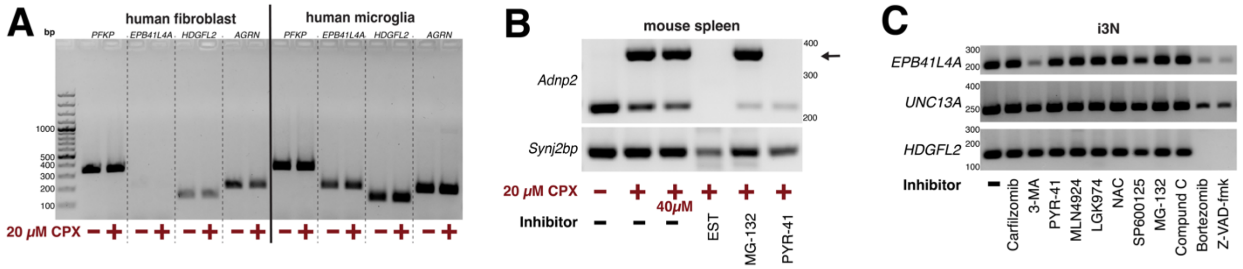


**Supplementary Figure 2: Further testing of CPX and inhibitors displays cell-type specific CPX response, along with differential response to inhibitors.** (**A**) Human fibroblasts and iPS-derived microglia treated with 20 μM CPX for four hours do not incorporate cryptic exons (**B**) Mouse spleen treated with various inhibitors in combination with CPX (**C**) i3N treated with inhibitors only (no CPX) do not incorporate cryptic exons after 4.5 hours. i3N treated with 3-MA, Z-VAD-fmk, and Bortezomib have reduced levels of RNA and viability.

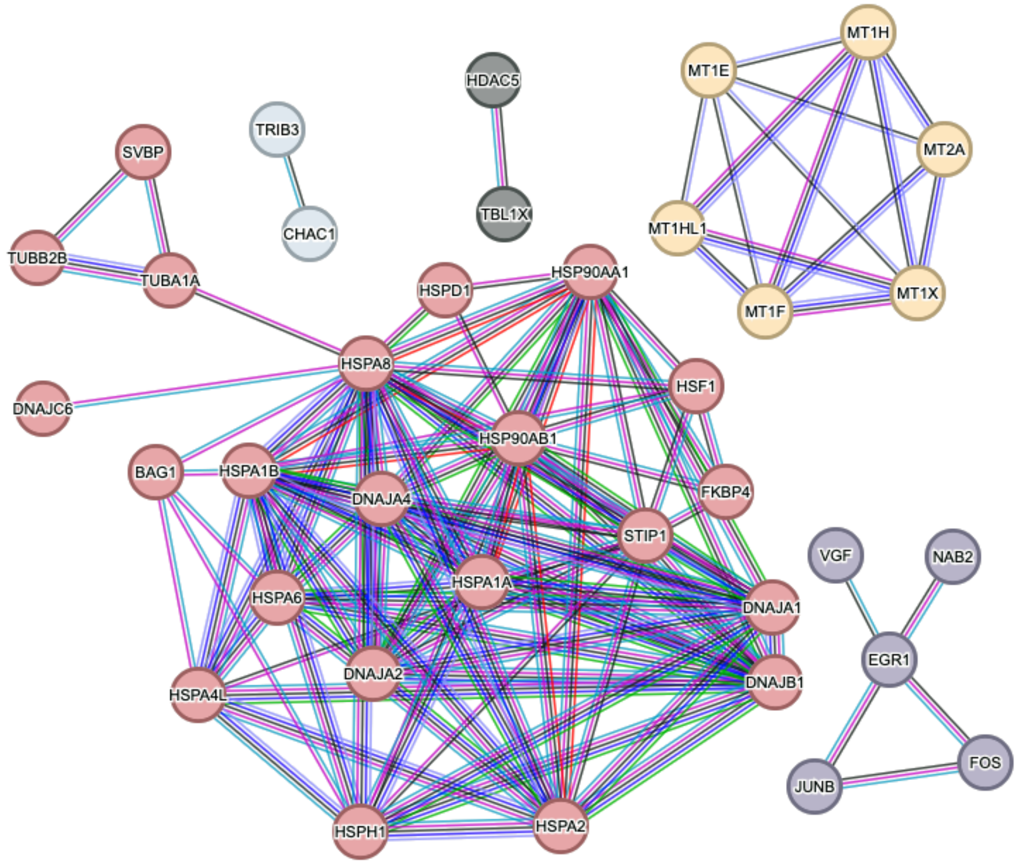


**Supplementary Figure 3: Network Analysis of Gene Interactions in i3Neurons Treated with CPX via STRING Database.** Protein-protein interaction networks from STRING show two main clusters among genes with $\log_2$(fold change) > 2 following CPX treatment in i3Neurons: metallothionein genes and heat-shock proteins, indicating responses to metal toxicity and cellular stress, respectively.

| Gene | Inclusion Junction 1 Coordinates | Inclusion Junction 2 Coordinates | Exclusion Junction Coordinates |
|---|---|---|---|
| ACTL6B | chr7:100650136-100650574 | chr7:100650644-100655018 | chr7:100650136-100655018 |
| AGRN_short | chr1:1044440-1044891 | chr1:1044988-1045160 | chr1:1044440-1045160 |
| AGRN_long | chr1:1044440-1044891 | chr1:1045081-1045160 | chr1:1044440-1045160 |
| ATG4B_long | chr2:241668686-241668781 | chr2:241668986-241670725 | chr2:241668686-241670725 |
| ATG4B_short | chr2:241668686-241668874 | chr2:241668986-241670725 | chr2:241668686-241670725 |
| EPB41L4A | chr5:112266331-112267209 | chr5:112267285-112275325 | chr5:112266331-112275325 |
| G3BP1 | chr5:151786716-151787764 | chr5:151787795-151790322 | chr5:151786716-151790322 |
| GPSM2 | chr1:108885579-108896316 | chr1:108896432-108896863 | chr1:108885579-108896863 |
| HDGFL2 | chr19:4491836-4492014 | chr19:4492153-4493702 | chr19:4491836-4493702 |
| PFKP | chr10:3099353-3099556 | chr10:3099820-3101364 | chr10:3099353-3101364 |
| RANBP1 | chr22:20122422-20122579 | chr22:20122698-20125307 | chr22:20122422-20125307 |
| STMN2 | chr8:79611215-79616821 | NA | chr8:79611215-79636801 |
| UNC13A | chr19:17641557-17642413 | chr19:17642542-17642844 | chr19:17641557-17642844 |
| UNC13A_alt | chr19:17641557-17642413 | chr19:17642592-17642844 | chr19:17641557-17642844 |
| UNC13B | chr9:35313990-35364544 | chr9:35364568-35366946 | chr9:35313990-35366946 |
| SLC24A3 | chr20:19681992-19683912 | chr20:19683961-19684175 | chr20:19681992-19684175 |
| IGLON5 | chr19:51311927-51320385 | chr19:51320671-51322063 | chr19:51311927-51322063 |
| KALRN_long | chr3:124700034-124700976 | chr3:124701256-124702037 | chr3:124700034-124702037 |
| KALRN_short | chr3:124700034-124701092 | chr3:124701256-124702037 | chr3:124700034-124702037 |
| MYO18A | chr17:29122254-29131382 | chr17:29131443-29165941 | chr17:29122254-29165941 |
| MYO18A_alt | chr17:29124689-29131382 | chr17:29131443-29165941 | chr17:29124689-29165941 |
| RSF1 | chr11:77764690-77813212 | chr11:77813784-77820527 | chr11:77764690-77820527 |
| SYT7 | chr11:61547309-61547529 | chr11:61547621-61551383 | chr11:61547309-61551383 |
| SYNE1 | chr6:152244657-152247822 | chr6:152247945-152249160 | chr6:152244657-152249160 |
| CAMK2B | chr7:44254608-44258489 | chr7:44258611-44258871 | chr7:44254608-44258871 |
| PXDN | chr2:1649676-1650799 | chr2:1650887-1653627 | chr2:1649676-1653627 |
| TRRAP | chr7:98881251-98881694 | chr7:98881737-98881974 | chr7:98881251-98881974 |

**Table 1: Coordinates of queried human cryptic exon junctions using hg38 assembly.**

| Gene | Inclusion Junction 1 Coordinates | Inclusion Junction 2 Coordinates | Exclusion Junction Coordinates |
|---|---|---|---|
| Adnp2 | chr18:80137574-80138152 | chr18:80138305-80142648 | chr18:80137574-80142648 |
| Bud23 | chr5:135061345-135063780 | NA | chr5:135061345-135063883 |
| Crem | chr18:3276730-3278080 | chr18:3278225-3280948 | chr18:3276730-3280948 |
| Fam135a | chr1:24022006-24022506 | NA | chr1:24022006-24024212 |
| Hdac4 | chr1:92054973-92141519 | chr1:92141582-92148255 | chr1:92054973-92148255 |
| Ift81 | chr5:122555543-122556905 | chr5:122556992-122559154 | chr5:122555543-122559154 |
| Pnpla6 | chr8:3524330-3524947 | NA | chr8:3524330-3530961 |
| Slc7a6 | chr8:106169257-106179101 | NA | chr8:106168955-106179101 |
| Smg5 | chr3:88336498-88340648 | chr3:88340764-88341857 | chr3:88336498-88341857 |
| Spata7 | chr12:98634320-98634590 | chr12:98634645-98637565 | chr12:98634320-98637565 |
| Synj2bp | chr12:81510052-81510828 | NA | chr12:81504640-81510828 |
| Tbc1d1 | chr5:64339732-64339816 | chr5:64339927-64345251 | chr5:64339732-64345251 |
| Tecpr1 | chr5:144214420-144215737 | chr5:144215791-144216252 | chr5:144214420-144216252 |
| Unc13a | chr8:71666671-71667328 | chr8:71667373-71671625 | chr8:71666671-71671625 |

**Table 2: Coordinates of queried mouse cryptic exon junctions using mm10 assembly.**
